# Chemogenetic actuator drugs impair prefrontal cortex-dependent working memory in rhesus monkeys

**DOI:** 10.1101/864140

**Authors:** Nicholas A. Upright, Mark G. Baxter

## Abstract

The most common chemogenetic neuromodulatory system, Designer Receptors Exclusively Activated by Designer Drugs (DREADDs), uses a non-endogenous actuator ligand to activate a modified muscarinic acetylcholine receptor that is no longer sensitive to acetylcholine. It is crucial in studies using these systems to test the potential effects of DREADD actuators prior to any DREADD transduction, so that effects of DREADDs can be attributed to the chemogenetic system rather than the actuator drug. We investigated working memory performance after injections of three DREADD agonists, clozapine, olanzapine, and deschloroclozapine, in male rhesus monkeys tested in a spatial delayed response task. Performance at 0.1 mg/kg clozapine and 0.1 mg/kg deschloroclozapine did not differ from mean performance after vehicle in any of the four subjects. Administration of 0.2 mg/kg clozapine impaired working memory function in three of the four monkeys. Two monkeys were impaired after administration of 0.1 mg/kg olanzapine and two monkeys were impaired after the 0.3 mg/kg dose of deschloroclozapine. We speculate that the unique neuropharmacology of prefrontal cortex function makes the primate prefrontal cortex especially vulnerable to off-target effects of DREADD actuator drugs with affinity for endogenous monoaminergic receptor systems. These findings underscore the importance of within-subject controls for DREADD actuator drugs to confirm that effects following DREADD receptor transduction are not due to the actuator drug itself, as well as validating the behavioral pharmacology of DREADD actuator drugs in the specific tasks under study.

**Significance Statement:** Chemogenetic technologies, such as Designer Receptors Exclusively Activated by Designer Drugs (DREADDs), allow for precise and remote manipulation of neuronal circuits. In the present study, we tested monkeys in a spatial delayed response task after injections of three actuator drugs – clozapine, olanzapine, and deschloroclozapine. We found that monkeys showed significant working memory impairments after 0.2 mg/kg clozapine, 0.1 mg/kg olanzapine, and 0.3 mg/kg deschloroclozapine compared to vehicle performance. In monkeys that showed impairments, these deficits were particularly apparent at longer delay periods. It is imperative to validate the drugs and dosages in the particular behavioral test to ensure any behavior after DREADD transduction can be attributed to activation of the receptors and not administration of the actuator drug itself.

## Introduction

Chemogenetic technologies are widely used to reversibly target specific neuronal populations and circuitry in awake, behaving animals (Eldridge et al., 2016; Grayson et al., 2016; Upright et al., 2018; Malvaez et al., 2019; Vetere et al., 2019). The most common chemogenetic system uses designer receptors exclusively activated by designer drugs (DREADDs) composed of a modified muscarinic acetylcholine receptor that is no longer sensitive to acetylcholine but is responsive to an otherwise inert drug (Armbruster et al., 2007). Clozapine-*N*-oxide (CNO), a metabolite of the antipsychotic clozapine, was initially the primary actuator drug used with DREADDs (Roth, 2016). However, drawbacks of CNO include poor brain penetrance, action as a substrate for P-glycoprotein, and reverse metabolism to its parent compound (Gomez et al., 2017; Raper et al., 2017; Mahler and Aston-Jones, 2018; Manvich et al., 2018). Together these shortcomings have sparked the need for new actuator drugs to use with DREADDs (Chen et al., 2015; Weston et al., 2018; Bonaventura et al., 2019; Goutaudier et al., 2019).

Three of these new actuators – low-dose clozapine, low-dose olanzapine, and deschloroclozapine – possess better brain penetrance and higher affinity and potency for DREADD receptors than CNO. All of these actuators have been tested in rodent systems (Gomez et al., 2017; Thompson et al., 2018; Weston et al., 2018) and low-dose clozapine (Raper et al., 2019) and deschloroclozapine (Nagai et al., 2019) have been tested in nonhuman primates. Clozapine and olanzapine are atypical antipsychotics designed for clinical use, and deschloroclozapine is a metabolite of clozapine; each has high affinity for DREADD receptors (Gomez et al., 2017; Weston et al., 2018; Nagai et al., 2019). Importantly, these drugs can activate DREADD receptors at doses lower than those associated with their therapeutic effects (Casey, 1993; Lidow and Goldman-Rakic, 1997; Murphy, 1997; Linn et al., 2003).

In this study, we tested two doses each of these three different DREADD actuators on a spatial working memory task in rhesus monkeys that had not yet received DREADD transduction. We tested all of these actuators so that we could select drugs to use as part of a subsequent study of DREADD neuromodulation in nonhuman primates. It is important in studies using any actuator with these chemogenetic systems to test the potential effects of the actuator prior to DREADD transduction in the behavioral task, so that effects of DREADDs can be attributed to the chemogenetic system and not the actuator drug itself.

## Materials and Methods

### Subjects

Four male rhesus macaques, noted as Cases A, J, Ro, and Ru, aged between 5 and 6 years and weighing 5.5 - 7.9 kg at the time of injection, were used for this study. Monkeys were socially housed indoors in single-sex groups. Daily meals consisted of monkey chow and a variety of fruits and vegetables, and were distributed within transport cages once testing was completed. On weekends, monkeys were fed in their home cages. Within the home cage, water was available *ad libitum*. Environmental enrichment, in the form of play objects and small food items, was provided daily in the home cage. All procedures were approved by the Icahn School of Medicine Institutional Animal Care and Use Committee and conform to NIH guidelines on the use of nonhuman primates in research.

### Apparatus

Testing was performed within a Wisconsin General Testing Apparatus (WGTA). The WGTA is a small enclosed testing area where the experimenter can manually interact with the monkey during testing. Monkeys were trained to move from the home cage to a metal transport cage which was then wheeled into the WGTA. The experimenter was hidden from the monkey’s view by a one-way mirror, with only the experimenter’s hands visible. A sliding tray with two food wells could be maneuvered back-and-forth between the experimenter and the monkey. A pulley-operated opaque black screen could be lowered by the experimenter to separate the tray from the monkey during testing.

### Behavioral Testing

Training on the delayed response task followed Bachevalier and Mishkin, 1986 and Croxson et al., 2011. Monkeys were first shown a small food reward (craisin or M&M) that was placed in one of two food wells on a sliding tray. The left/right location of the reward was chosen across trials based on a pseudorandomly, counterbalanced sequence. Both wells were then covered with flat, gray tiles and a black opaque screen was lowered between the tray and the monkey for a predefined delay period. The screen was subsequently raised, and the test tray was advanced to the monkey, allowing the monkey to displace one of the well covers (**Figure 1**). During initial shaping, monkeys were taught to displace the tiles covering the wells and select the reward. Once monkeys readily displaced the tiles, they progressed through three stages of training with 24 trials per session. Once a monkey reached criterion on the third stage of training, experimental training began. In the experimental task, each trial consisted of one of five possible delays (5, 10, 15, 20, or 30 s) and left or right well location. These delay/location pairings were varied pseudorandomly across trials such that each delay occurred six times and each well location occurred fifteen times in each testing session. Monkeys continued with the variable delay paradigm for the remainder of the experiment. Drug injections began once a monkey had achieved stable performance on the task.

**Figure 1.**
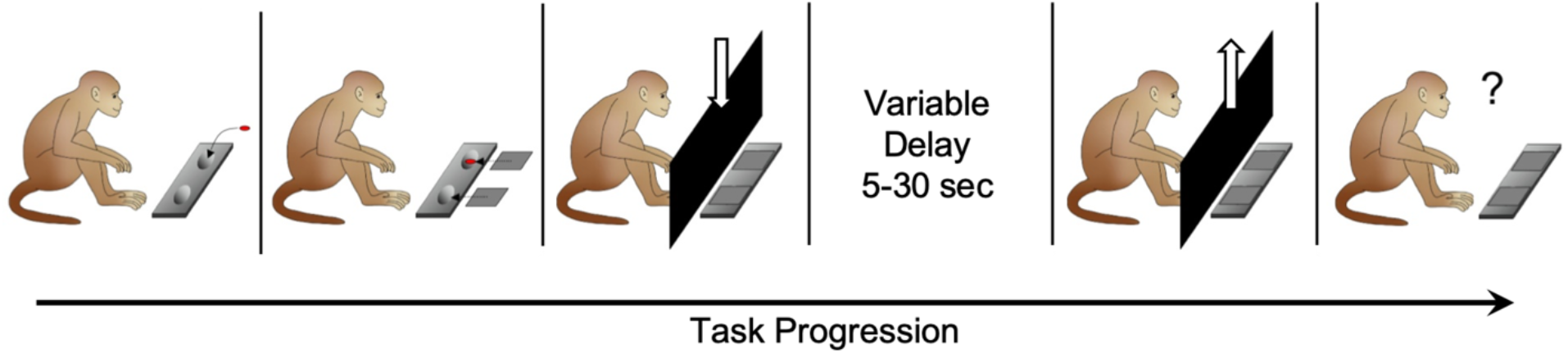
Spatial delayed response task. In the first sample phase, the monkey views a reward placed into one of two wells. The wells are covered and hidden from the monkey by an opaque screen for a variable delay, ranging between 5-30 seconds. The screen is raised, and the monkey must select the well containing the reward.

### Drugs

#### Pharmacology

Structures for clozapine, olanzapine, and deschloroclozapine can be found in **Figure 2**. Clozapine, marketed as the prototype atypical antipsychotic Clozaril^TM^, is FDA-approved and treats psychosis related to schizophrenia and Parkinson’s disease (Tauscher et al., 2004; Zhang et al., 2019). It is a dopamine receptor antagonist with an affinity for D2 and D4 receptors and shows high affinity for serotonin 5-HT2A receptors (Tauscher et al., 2004; Wenthur and Lindsley, 2013). A dose of 0.1 mg/kg clozapine is effective at activating DREADD receptors in vivo in monkeys (Raper et al., 2019). Olanzapine, a second-generation atypical antipsychotic and also FDA-approved, has been documented as a potentially safer, yet not as effective, alternative to clozapine. Similar to clozapine, olanzapine displays antagonism primarily for dopamine D2 and serotonin 5-HT2A receptors (Lidow and Goldman-Rakic, 1997; Tauscher et al., 2004). It has recently been shown that olanzapine is a potent agonist at the inhibitory hM4D(Gi) DREADD receptor in vivo (Weston et al., 2018). Both actuators are FDA-approved which could facilitate their application for chemogenetic technologies in a clinical setting. Deschloroclozapine is a dual dopamine and serotonin receptor antagonist with some selectivity for D1, 5-HT2A, and 5-HT2C receptors (Nagai et al., 2019).

**Figure 2.**
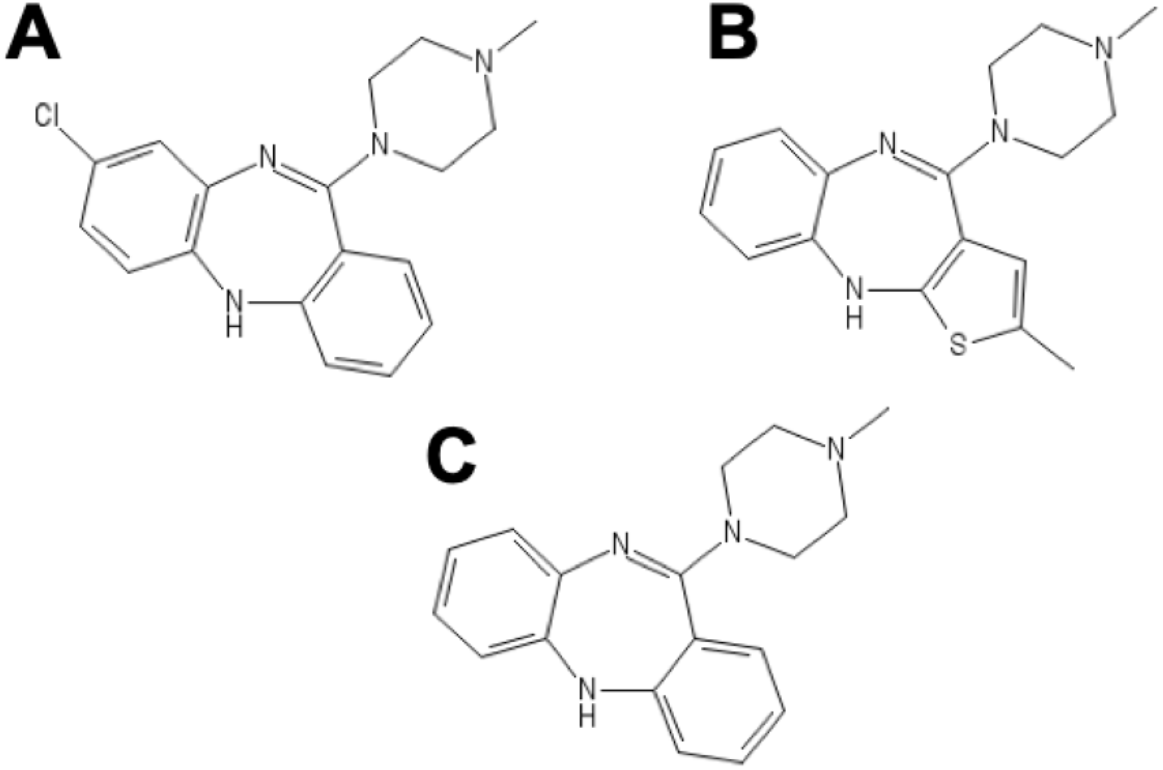
Chemical structures of DREADD actuator drugs. **A,** clozapine. **B,** olanzapine. **C,** deschloroclozapine.

#### Preparation

Drugs were prepared fresh daily, at concentrations so that monkeys received 0.1 ml/kg for injection (e.g., for a 0.2 mg/kg dose of drug, drug solution was prepared at a concentration of 2.0 mg/ml). Solutions were filtered through a 0.22 um syringe filter and pH was determined before injection. Acetic acid, sodium acetate, and sodium hydroxide (NaOH) were all obtained from Fisher Scientific. Concentrations used for glacial acetic acid, sodium acetate, and NaOH were 99.7% (v/v), 1 M, and 0.2 M respectively. Clozapine (Tocris, Minneapolis, MN) was stored at room temperature. Clozapine was given at 0.1 or 0.2 mg/kg, intramuscularly. For 0.1 mg/kg, clozapine powder was first dissolved in acetic acid and sodium acetate then diluted with NaOH to a final concentration in .25/50/49.75 acetic acid/sodium acetate/NaOH (v/v/v). For 0.2 mg/kg, the same reagents were used but the final concentration was 0.5/50/49.5 acetic acid/sodium acetate/NaOH (v/v/v). Olanzapine (Tocris, Minneapolis, MN) was stored at room temperature and given at 0.05 or 0.1 mg/kg. Olanzapine solutions were made using the same method as above for the 0.1 mg/kg clozapine dose. Deschloroclozapine (DCZ; Medchemexpress, Monmouth Junction, NJ) was stored at 4 °C and was given at 0.1 mg/kg and 0.3 mg/kg. Low dose of deschloroclozapine was made using the same method as low-dose clozapine and high dose deschloroclozapine was made using the same method as the higher dose of clozapine. Vehicle injections consisted of .25/50/49.75 acetic acid, sodium acetate, and NaOH and were given at 0.1 ml/kg. Actuator injections were never given more than twice in one week, with vehicle or no-injection test days on other days of the test week. Clozapine and olanzapine were given in the home cage 10 min before the start of testing and deschloroclozapine was given in the home cage 30 min before the start of testing.

### Statistical Analysis

Analyses were performed in RStudio with R 3.5 (R Core Team, 2018) using the lme4, lmerTest, and emmeans packages (Kuznetsova et al., 2017; Lenth, 2019). Data were analyzed with a generalized linear mixed model and binomial distribution, with trial outcome (correct or incorrect) as the outcome variable, Injection and Delay as fixed effects, and Case as a random effect to account for repeated sessions across time for each monkey. Two models were estimated for each drug condition (CLZ, OLZ, and DCZ) across all four cases comparing the two drug doses in each condition to vehicle (VEH). A “full” model included a two-way interaction term between Injection (VEH, 0.1 mg/kg CLZ, 0.2 mg/kg CLZ for clozapine; VEH, 0.05 mg/kg OLZ, 0.1 mg/kg OLZ for olanzapine; VEH, 0.1 mg/kg DCZ, 0.3 mg/kg DCZ for deschloroclozapine) and Delay (5-30 seconds) to examine how actuator injection affected working memory performance over delay intervals during the delayed response task. A “reduced” model with Delay alone as a fixed effect was estimated and, using a hierarchical approach, we assessed model fit between the two models. If Injection significantly improved model fit (i.e., there was a significant effect of injection on task performance) we calculated estimated marginal means from the “full” fitted model for each drug condition across all four cases. We analyzed pairwise comparisons and contrasts to determine differences in performance between VEH and each actuator drug dose as well as differences between the two doses of each actuator drug themselves. Due to inter-case variability across monkeys in response to actuator drug, a generalized linear model for each drug condition was also computed for each Case regardless of the outcome of the overall model. We next calculated estimated marginal means from the “full” fitted model for each Case and analyzed pairwise contrasts to determine any differences in performance between 1) VEH and each actuator drug dose and 2) two doses of each actuator drug themselves. Interaction analyses were calculated using the estimated marginal means to examine whether a dose of an actuator drug modified the working performance over delay for each monkey. Reported confidence intervals are given on a log odds ratio scale. P-values and confidence intervals were adjusted using the Tukey method for multiple comparisons. Because three out of four monkeys had no variability at a particular delay or delays during actuator drug sessions, the five levels for the Delay factor were collapsed into “short delay” and “long delay” levels for some analyses (noted for each monkey). Full models, those for each drug condition across all four cases and those for individual cases, were tested for over/underdispersion of residuals using simulation-based tests to measure deviation of residuals and dispersion of residual standard deviations (Hartig, 2019). All evaluations of model residuals returned no significant over/underdispersion (two-sided nonparametric test, p > 0.05).

## Results

### Effect of clozapine on working memory performance

Comparison of our full and reduced models showed that including Injection and Injection x Delay interaction terms significantly improved our model fit (*X*^2^(10) = 32.209, p = 0.0003695). Analysis of the estimated marginal means comparisons from our full model across all four cases revealed that monkeys showed significant working memory impairment after 0.2 mg/kg clozapine compared to vehicle (**Figure 3A**; p = 0.0007, 95% confidence interval of [0.173, 0.779]). We found a significant effect on working memory performance between the two doses of clozapine (p = 0.001, [0.222, 1.058]). Pairwise comparisons of each vehicle-clozapine dose pair across all four monkeys revealed a significant effect of the higher 0.2 mg/kg clozapine dose relative to vehicle and 0.1 mg/kg clozapine at the 30 second delay interval only (p = 0.0002). For the analysis of pairwise contrasts for each case, delays were binned into “short” (5-20 s) and “long” (30 s) groups. Comparisons of estimated marginal means for each monkey showed that Cases A, Ro, and Ru were significantly impaired in the spatial delayed response task after 0.2 mg/kg clozapine compared to performance after vehicle (**Table 1**). Interestingly, Case A also showed a significant difference in working memory performance between the two doses of clozapine. Case A had a significant interaction between Injection and Delay terms (p = 0.0103). This interaction was primarily driven by the longer delay period with worsening delayed response performance with the higher clozapine dose and the long 30s delay group (p = 0.0005). No monkey showed impairment in working memory performance after the 0.1 mg/kg clozapine dose compared to vehicle performance.

**Figure 3.**
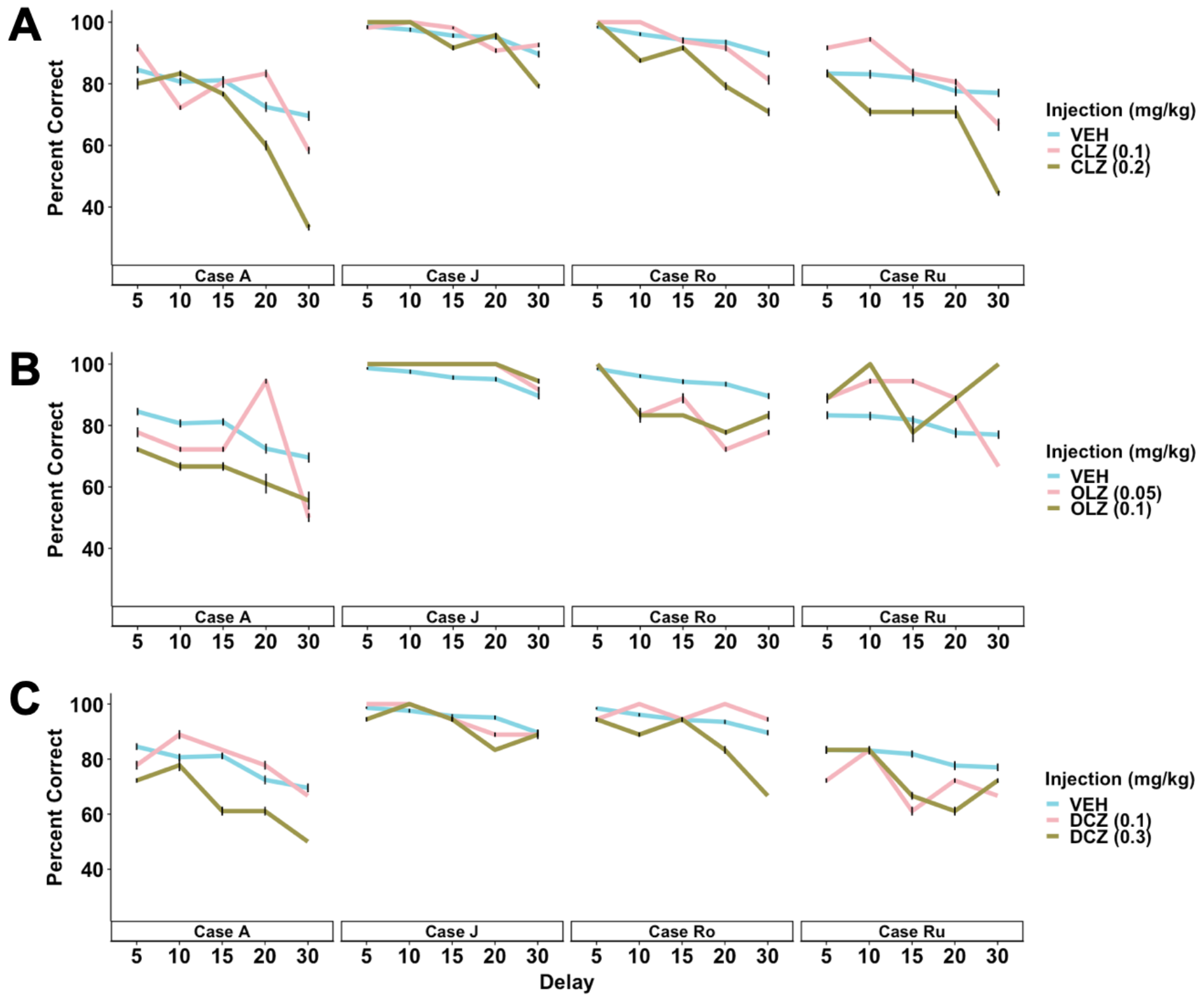
Spatial delayed response performance for vehicle and each drug condition. **A,** Performance after clozapine injection for each case across all tested delay intervals. **B,** Performance after administration of olanzapine. **C,** Performance after administration of deschloroclozapine. Data are represented as mean performance ± sem. VEH, vehicle; CLZ, clozapine; OLZ, olanzapine; DCZ, deschloroclozapine.

**Table 1.**
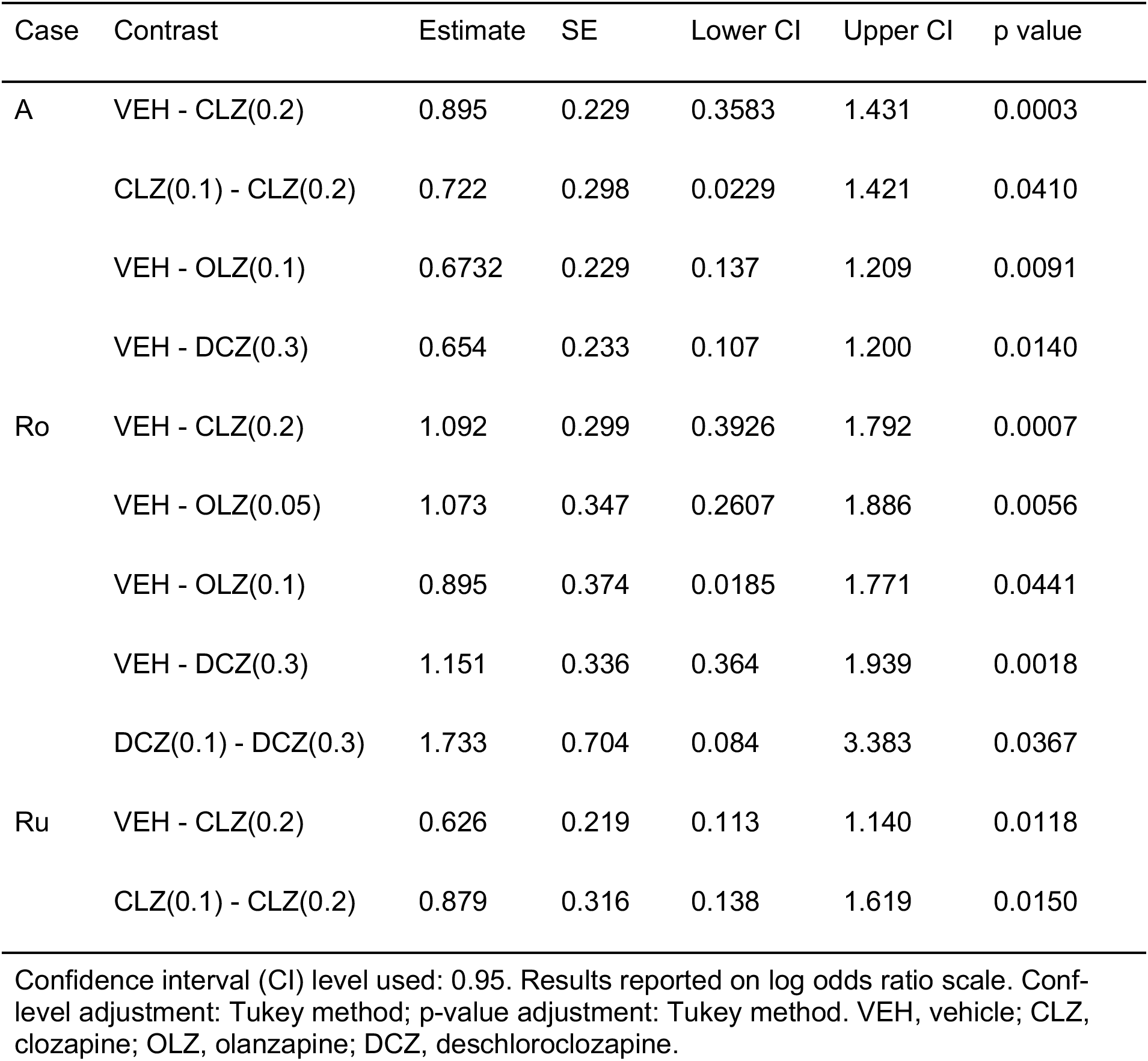
Significant contrasts of estimated marginal means by Case

| Case | Contrast | Estimate | SE | Lower CI | Upper CI | p value |
| --- | --- | --- | --- | --- | --- | --- |
| A | VEH - CLZ(0.2) | 0.895 | 0.229 | 0.3583 | 1.431 | 0.0003 |
|  | CLZ(0.1) - CLZ(0.2) | 0.722 | 0.298 | 0.0229 | 1.421 | 0.0410 |
|  | VEH - OLZ(0.1) | 0.6732 | 0.229 | 0.137 | 1.209 | 0.0091 |
|  | VEH - DCZ(0.3) | 0.654 | 0.233 | 0.107 | 1.200 | 0.0140 |
| Ro | VEH - CLZ(0.2) | 1.092 | 0.299 | 0.3926 | 1.792 | 0.0007 |
|  | VEH - OLZ(0.05) | 1.073 | 0.347 | 0.2607 | 1.886 | 0.0056 |
|  | VEH - OLZ(0.1) | 0.895 | 0.374 | 0.0185 | 1.771 | 0.0441 |
|  | VEH - DCZ(0.3) | 1.151 | 0.336 | 0.364 | 1.939 | 0.0018 |
|  | DCZ(0.1) - DCZ(0.3) | 1.733 | 0.704 | 0.084 | 3.383 | 0.0367 |
| Ru | VEH - CLZ(0.2) | 0.626 | 0.219 | 0.113 | 1.140 | 0.0118 |
|  | CLZ(0.1) - CLZ(0.2) | 0.879 | 0.316 | 0.138 | 1.619 | 0.0150 |

Confidence interval (CI) level used: 0.95. Results reported on log odds ratio scale. Conf-level adjustment: Tukey method; p-value adjustment: Tukey method. VEH, vehicle; CLZ, clozapine; OLZ, olanzapine; DCZ, deschloroclozapine.

### Effect of olanzapine on working memory performance

Analysis of our full and reduced models revealed that inclusion of Injection and Injection x Delay terms did not significantly improve model fit (*X*^2^(10) = 6.558, p = 0.7664) indicating there was no consistent difference across the four cases. To examine variability within our cases, we also computed a full model for each monkey. For Case Ro, delays were binned into the same “short” (5-20s) and “long” (30s) groups. We found that Cases A and Ro were significantly impaired in working memory performance after administration of the higher 0.1 mg/kg dose of olanzapine (**Figure 3B**). Case Ro also had significantly more incorrect trials in the delayed response task after the lower 0.05 mg/kg dose of olanzapine compared to vehicle. Although not quantified, it was apparent during testing that Cases A, J, and Ro displayed an increased amount of lachrymation and cooing behaviors during each olanzapine testing session; no noticeable increase in these behaviors was noted for vehicle or other actuator drugs.

### Effect of deschloroclozapine on working memory performance

Including Injection and Injection x Delay interaction terms significantly improved our deschloroclozapine model compared to Delay alone (*X*^2^(10) = 19.686, p = 0.03237).

Comparison of estimated marginal means for our full fitted model across all four cases revealed a significant contrast between vehicle and the higher 0.3 mg/kg dose of deschloroclozapine (p = 0.0058, 95% confidence interval of [0.1093, 0.802]). Analysis of the estimated marginal means for the full model for each case showed Cases A and Ro were significantly impaired in delayed response performance after 0.3 mg/kg dose compared to performance after vehicle (**Table 1**). Due to no variability at two delays, Case Ro again had binned “short” and “long” delays. Case Ro also had a significant difference between the two doses of deschloroclozapine with greater working memory deficit after the 0.3 mg/kg dose. No monkey showed a significant impairment in working memory performance after the lower 0.1 mg/kg dose of DCZ (**Figure 3C**).

## Discussion

We found that some monkeys showed deficits in the delayed response task after administration of actuator drugs prior to any DREADD transduction. For clozapine, three out of four monkeys were impaired in working memory function by the higher 0.2 mg/kg dose which is far less than doses tested for therapeutic effects (ranging 2.5-6 mg/kg; Casey, 1993; Lidow and Goldman-Rakic, 1997; Murphy, 1997; Linn et al., 2003). Two out of four monkeys showed working memory deficits after the higher 0.1 mg/kg dose of olanzapine and one of those monkeys, Case Ro, additionally showed significant impairment compared to vehicle after the 0.05 mg/kg dose. Similar to clozapine, both of the tested doses for olanzapine are below a therapeutic dose of 0.35 mg/kg yet have been shown to effectively activate DREADDs (Lidow and Goldman-Rakic,1997; Weston et al., 2018). The two monkeys impaired by doses of olanzapine were also impaired by the higher 0.3 mg/kg dose of deschloroclozapine. We found that no monkey was impaired in delayed response performance after 0.1 mg/kg clozapine or 0.1 mg/kg deschloroclozapine and we plan on moving forward with those actuators and doses for our future neuromodulatory studies.

We speculate that the delayed response task may be particularly sensitive to the actuators used in this study due to the neuropharmacology of the prefrontal cortex, a region highly implicated in working memory function (Goldman-Rakic, 1995; Fuster, 2001; Arnsten et al., 2012; Barbey et al., 2013). Indeed, dopamine and serotonin receptor subtypes antagonized by these three actuator drugs are enriched in primate prefrontal cortex, and D2 and 5-HT2A specifically have been implicated in working memory function (Puig and Gulledge, 2011; Ott and Nieder, 2016). We found a slight dependence on delay with the higher doses of clozapine and deschloroclozapine on delayed response performance with greater impairment at higher delay intervals (20-30 s). This behavioral deficit at higher delay periods would point to a working memory effect and demonstrate that antagonism of these receptors did not impair delayed response performance overall, but instead performance where memory of reward location in the task must be maintained for the longest periods. Our goal for this study was not to examine the receptor mechanisms of these drugs, and in the absence of pharmacokinetic data on each monkey we do not know whether individual differences in drug response were related to individual differences in drug distribution and/or metabolism.

A recent study also investigated the role of acute systemic low-dose clozapine on a battery of behaviors in wild-type rats (Ilg et al., 2018). The authors found that low-dose clozapine (0.1 mg/kg) did not affect working memory function measured as performance in a delayed alternation task. The authors did find effects of clozapine on other behaviors such as locomotion, anxiety, and cognitive flexibility. Thus, similar to our findings in nonhuman primates, they found that actuator drugs can have DREADD-independent effects at a high enough dose. We stress that our findings demonstrate that these behavioral effects can occur in a cognitive task with doses far below the therapeutic range tested in nonhuman primates (clozapine: 6 mg/kg; olanzapine: 0.35 mg/kg) that would normally not be expected to have any effects on their own.

We selected these three actuator drugs due to their affinity and efficacy at DREADD receptor and their evidence in rodent and nonhuman primate behavioral models. Two recently developed compounds, JHU37152 (J52) and JHU37160 (J60), also display high in vivo occupancy and potency for DREADD receptors in mice and monkeys (Bonaventura et al., 2019). These drugs exhibit greater DREADD activation than clozapine at lower concentrations, indicating that they are more potent agonists at DREADD receptors. In contrast, another actuator drug, Compound 21, exhibits similar potency to CNO at the DREADD receptor; however, this potency is lower than both J52 and J60 and thus may not be practical for applications in nonhuman primates (Chen et al., 2015; Thompson et al., 2018; Bonaventura et al., 2019). Recently, it was reported that low-dose Compound 21 also produced metabolic changes in brain activity (FDG uptake) while an equipotent dose of clozapine produced no changes (Bonaventura et al., 2019). Administration of 0.1 mg/kg clozapine and 0.1 mg/kg deschloroclozapine led to no significant working memory impairment in any monkey and we plan on using these actuators moving forward; however we recognize these alternatives for effective actuator drugs, particularly J52 and J60, and could test these on cognitive behaviors in future neuromodulatory studies.

While the most commonly used form of DREADDs uses the modified G-protein coupled receptor, other chemogenetic technologies, including the KORD DREADD derived from a modified k-opioid receptor (Vardy et al., 2015) and the recently reported pharmacologically selective actuator modules (PSAMs; Magnus et al., 2019), are available to modulate neuronal regions and circuitry. For our study, we selected the muscarinic acetylcholine receptor-based DREADDs due to its extensive application in nonhuman primates both in our own behavioral studies as well as several others (Eldridge et al., 2016; Grayson et al., 2016; Upright et al., 2018; Raper et al., 2019). There has been limited evidence of PSAMs in nonhuman primates and the low solubility of Salvanorin B, the actuator for KORD DREADD, makes dosing monkeys difficult. Any actuator drug or technology employed should be held to the same scrutiny as those in this study and researchers should plan similar preoperative experiments for any planned doses.

DREADDs remain a powerful tool remotely to manipulate neuronal activity in vivo. In this study, we examined the effects of three actuator drugs on working memory performance in four monkeys prior to DREADD transduction and isolated two drugs, each at 0.1 mg/kg dose, that did not cause any significant deficits in preoperative spatial delayed response performance. The presence of off-target behavioral effects of DREADD actuator drugs in this study may be related to the specific behavioral paradigm used. Higher doses of actuator drugs will increase occupancy at DREADD receptors and provide a greater chemogenetic effect, but will also increase the likelihood of off-target effects at endogenous receptors. For studies using DREADDs, particularly those in nonhuman primates, it would be critical to obtain an upper limit of an actuator dose for each animal prior to DREADD transduction and determine individual variability with dosages. After DREADDs are expressed, it is difficult, and in some cases not possible, to ascertain whether a “low” dose is effective on its own. Our findings also indicate that different potential DREADD actuators, even at “low” doses, may have different patterns of off-target effects in different behavioral paradigms, as in the case of olanzapine here. Therefore, we stress the importance of validating each drug and dosage in the context of the planned behavioral testing post-DREADD transduction and in a within-subjects design when possible.

Including these experiments in chemogenetic studies will ensure that any behavioral output post transduction can be attributed to the activated chemogenetic system and not administration of the actuator drug itself.

## Acknowledgements

We thank Jared Boyce for technical assistance. This work was supported by R21NS096936 (MGB) and T32AG049688 (NAU).

